# Transcriptome analysis of SARS-CoV-2 naïve and recovered individuals vaccinated with inactivated vaccine

**DOI:** 10.1101/2021.11.05.467537

**Authors:** Yuwei Zhang, Xingyu Guo, Cunbao Li, Zengqiang Kou, Lanfang Lin, Mingxiao Yao, Bo Pang, Xiaomei Zhang, Qing Duan, Xueying Tian, Yufang Xing, Xiaolin Jiang

## Abstract

The urgent approval of the use of the inactivated COVID-19 vaccine is essential to reduce the threat and burden of the epidemic on global public health, however, our current understanding of the host immune response to inactivated vaccine remains limited. Herein, we performed serum IgG antibody detection and transcriptomics analysis on 20 SARS-CoV-2 naïve individuals who received multiple doses of inactivated vaccine and 5 SARS-CoV-2 recovered individuals who received single dose of inactivated vaccine. Our research revealed the important role of many innate immune pathways after vaccination, identified a significant correlation with the third dose of booster vaccine and proteasome-related genes, and found that SARS-CoV-2 recovered individuals can produces a strong immune response to a single dose of inactivated vaccine. These results help us understand the reaction mechanism of the host’s molecular immune system to the inactivated vaccine, and provide a basis for the choice of vaccination strategy.

## Introduction

Since December 2019, a new severe acute respiratory syndrome coronavirus 2 (SARS-CoV-2) has swept the world, causing a variety of clinical syndromes termed coronavirus disease 2019 (COVID-19)(1). The clinical manifestations of COVID-19 include fever, dry cough, fatigue, sore throat, pneumonia, diarrhea and other symptoms, and may even develop into severe pneumonia, acute respiratory distress syndrome (ARDS) or multiple organ failure(2). The World Health Organization declared a pandemic in March 2020. COVID-19 has caused considerable impacts on the global economy and public health.

Although for a long time, people have relied on social distancing, hygiene measures, and repurposed drugs to coping with it, now many researchers are committed to developing safe and effective vaccines to establish herd immunity to prevent SARS-CoV-2. In view of the turmoil caused by the COVID-19 pandemic and the urgent need for effective vaccine, vaccine development can be accelerated by combining originally requested phases. The vaccine does not go through a complete approval process, but may be approved for emergency use(3). Early clinical trial results of the inactivated vaccines produced by Sinopharm and Sinovac showed a low incidence of adverse reactions and good immunogenicity(4–8). On July 22, 2020, the above two candidate inactivated vaccines were approved for use(9). These two vaccines are widely promoted and vaccinated, but the reaction mechanism of the host’s molecular immune response to the inactivated vaccine is not yet fully understood, and the implementation of the third booster dose is also being actively discussed recently(10) (https://www.cdc.gov/coronavirus/2019-ncov/vaccines/booster-shot.html). In addition, the impact of prior SARS-CoV-2 infection status on vaccination response is also worthy of further analysis. These insights may provide a theoretical basis for the determination of vaccination strategies and the allocation of vaccine resources.

The application of high-throughput technology to systematically scan the transcriptome response and evaluate changes in gene expression levels is very suitable for identifying immune response dynamics and gene regulatory networks. Previously, transcriptome analysis of Hantavax vaccine(11), influenza vaccine(12), VSV-EBOV vaccine(13) and BNT162b mRNA vaccine(14) have fully revealed the dynamics of the host immune response after vaccination. In this study, we characterized the PBMC transcriptome changes of SARS-CoV-2 recovered individuals receiving one dose of vaccine and healthy SARS-CoV-2 naïve individuals receiving one to three doses of vaccine respectively. This real-world study showed the changes of various cytokines and the regulation of immune pathways induced after vaccination, reveal the indispensable role of innate immune pathways, and reflect the key modules information of vaccine response in different individuals and different doses of vaccine.

## Methods

### Study population and recruitment

From January 2021, through August 2021, we recruited five SARS-CoV-2 recovered and twenty healthy SARS-CoV-2-naïve individuals to participate in this study in Linyi City, Shandong Province. Under different doses of the COVID-19 inactivated vaccine, anticoagulant and procoagulant blood samples were collected from participants. This study was approved by the Ethical Approval Committee of Shandong Center for Disease Control and Prevention.

### Serum and PMBCs isolation

The venous blood was collected from each participant to separate serum or isolate PBMCs. We separated sera by centrifugation at 2500 rpm/min for ten minutes and preserved at −80 °C until testing. PBMCs were isolated by density-gradient sedimentation using Lymphoprep™ density gradients (Axis-Shield, Norway), frozen in cell saving media and stored in liquid nitrogen.

### Qualitative SARS-CoV-2 IgG detection

The IgG antibodies were detected using an indirect ELISA kit (Beijing Wantai Biological Pharmacy Enterprise Co, China)(15, 16) based on a recombinant nucleoprotein of SARS-CoV-2. The specific operation is carried out in strict accordance with the instructions. The cut-off value for IgG is the mean OD value of three negative controls (if the mean absorbance value for three negative calibrators is < 0.03, take it as 0.03) + 0.16. A serum sample with an OD value ≥cut-off OD value was considered to be an anti-N IgG antibody positive.

### RNA-seq

Total RNA was extracted from PBMCs by using the RNeasy Mini Kit (Qiagen, Germany) according to the manufacturer’s instructions. The concentration and integrity of total RNA were checked using the Qubit RNA Assay Kit in Qubit 4.0 Fluorometer (Life Technologies, USA) and the RNA Nano 6000 Assay Kit of the Bioanalyzer 2100 System (Agilent Technologies, USA) respectively. A total amount of 100 ng total RNA per sample was used to prepare for the rRNA-depleted cDNA library by Stranded Total RNA Prep Ligation with Ribo-Zero Plus Kit (Illumina, USA). The final library size and quality was evaluated using an Agilent High Sensitivity DNA Kit (Agilent Technologies, CA), and the fragments were found to be between 250 and 350 bp in size. The library was sequenced using an Illumina NextSeq 2000 platform to generate 100 bp paired-end reads.

### Differentially expressed genes (DEGs) identification

RNA-seq data were processed quality control (QC), trimming and mapping to the human reference genome hg38 using CLC Genomics Workbench. Gene expression level was measured based on the transcripts per million (TPM). We calculated normalization factors using iterative edgeR(17) and limma(18) package, and filtered DEGs based on a p-value < 0.05 and a 2^logFC_cutoff. DEGs are visualized in the form of heat maps through the pheatmap (https://CRAN.R-project.org/package=pheatmap) package in R software.

### PPI network construction

The initial PPI network for the protein products of identified up and down regulated DEGs was constructed using the STRING Database(19) (STRING v11.5; https://string-db.org/), and then the network was visualized and analyzed with Cytoscape software(20). The CentiScape and MCODE plugins in the Cytoscape software were used to extract the characteristic genes from the DEGs.

### Pathway enrichment analysis

To be aware of the prospective functions of characteristic genes identified by the PPI network analysis, Kyoto Encyclopedia of Genes and Genomes (KEGG) pathways were identified using CluGO from Cytoscape software. P<0.05, subjected to Bonferroni adjustment, were defined as the cut-off criterion.

### Weighted Gene Co-expression Network Analysis (WGCNA)

We constructed the co-expression network through WGCNA package(21) in R software. The best β value was confirmed with a scale-free fit index bigger than 0.85 as well as the highest mean connectivity by performing a gradient test from 1 to 30. Subsequently, we transformed the adjacency matrix into a topological overlap matrix (TOM). The TOM obtained was then clustered by dissimilarity between genes, and we performed hierarchical clustering to identify modules, each containing at least 30 genes (minModuleSize=30). Some modules were combined according to the correlation coefficient. We calculated the correlation between the modules and the sample characteristics to identify significant modules. The genes in significant modules were analyzed using cytoHubba and GeneMANIA in Cytoscape software.

## Results

### Characteristic of samples

The median age of twenty-five participants was 40 years (interquartile range [IQR], 24–61), of which twelve (48%) were female and thirteen (52%) were male. They are divided into five groups according to their previous SARS-CoV-2 infection status and the difference in the number of doses of the COVID-19 inactivated vaccine. The Healthy-unvaccinated (H-un) group contains SARS-CoV-2 naïve individuals who have not been vaccinated with an inactivated vaccine; the Patient-1st (P-1st) group consists of COVID-19 patients who have received a single dose of vaccine and have recovered for 18 months; the Healthy-1st (H-1st) group includes SARS-CoV-2 naïve individuals who have received one dose of inactivated vaccine; the Healthy-2nd (H-2nd) group consists of SARS-CoV-2 naïve individuals who have received two doses of inactivated vaccine and the Healthy-3rd (H-3rd) group is SARS-CoV-2 naïve individuals who received the third booster vaccine half a year after receiving two doses of inactivated vaccine. We collected anticoagulant and procoagulant blood samples of participants fourteen days after vaccination. The specific details are shown in Fig. 1.

**Fig. 1.**
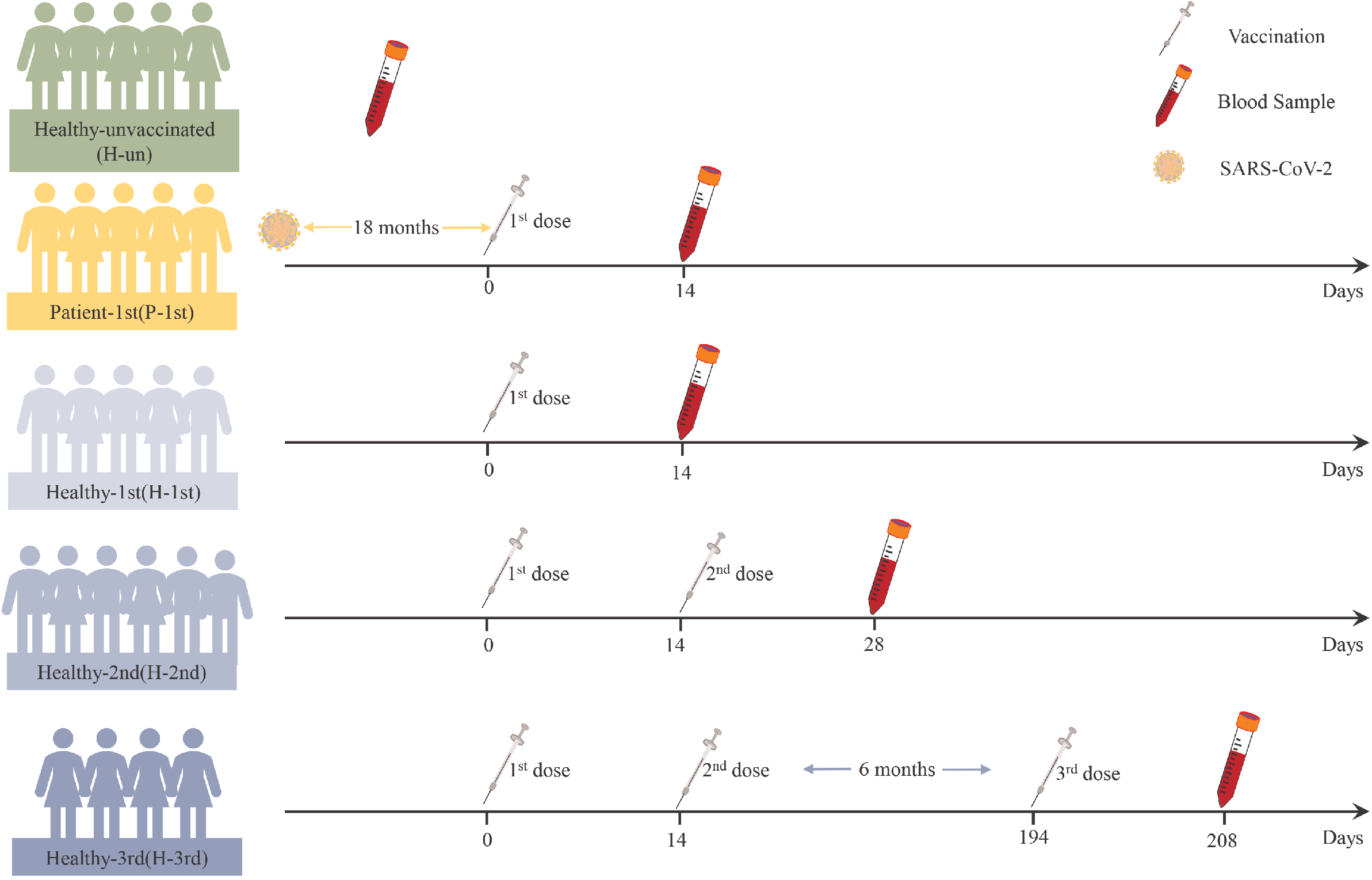
The details of this study design. Participants’ previous SARS-CoV-2 infection status, vaccine dose information and blood sample time points are shown.

### Detection of SARS-CoV-2 IgG in serum samples

In order to study the antibody response of serum samples to SARS-CoV-2 after vaccination, we used a fully validated commercial diagnostic ELISA kit to qualitatively measure IgG antibodies against N in serum samples diluted 1:10. The critical value for judging that the antibody level is positive is OD>0.19. The IgG antibodies of unvaccinated healthy participants were all negative, and only one healthy participant who received one dose of the vaccine was positive, whereas the IgG antibody results of recovered patients who received one dose of vaccine and healthy participants who received two or three doses of vaccine were both positive (Fig. 2).

**Fig. 2.**
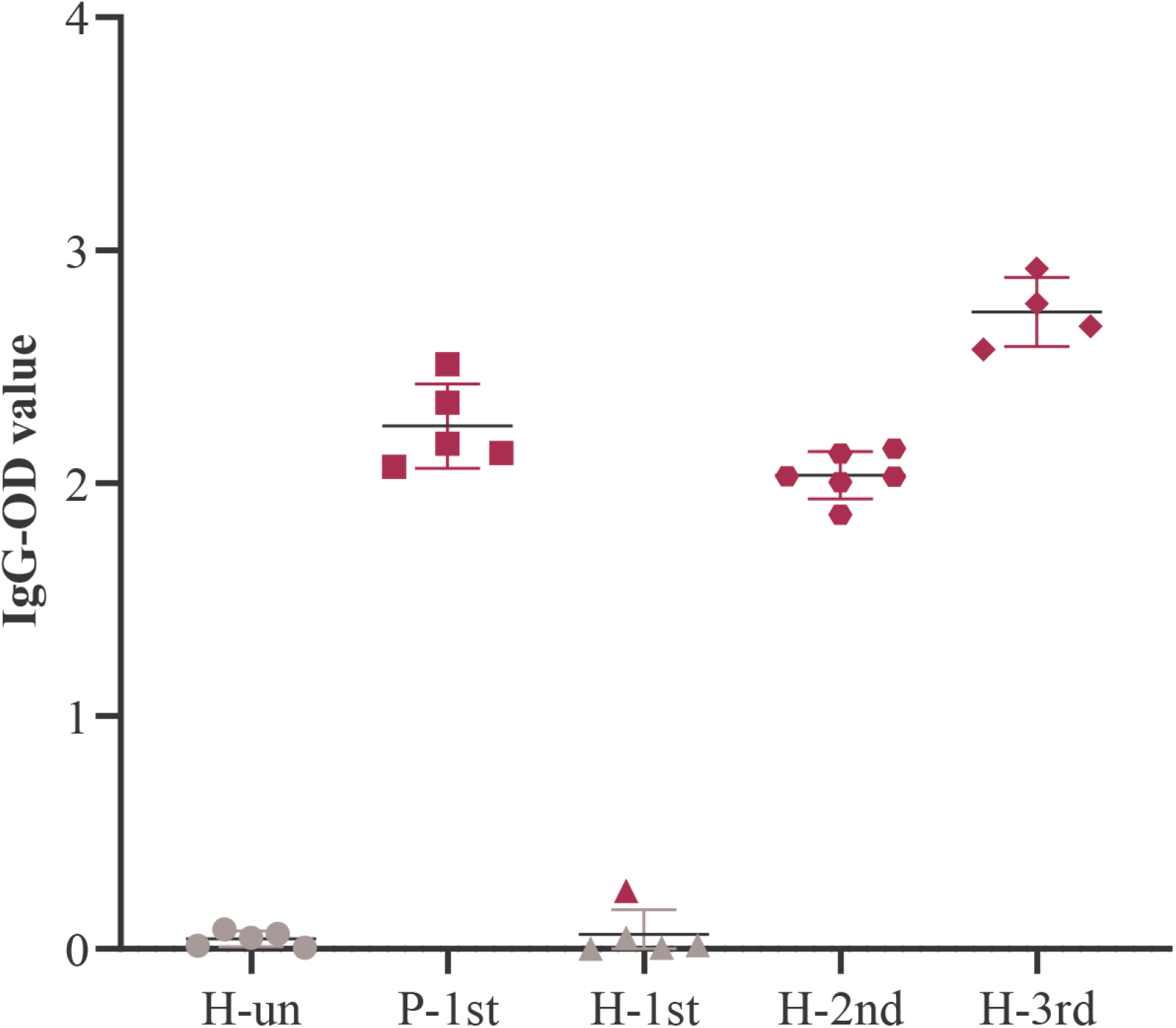
The level of IgG antibodies in the five groups of serum samples. Red represents positive and gray represents negative.

### Identification of DEGs

The transcriptome data of five groups of PBMCs under different vaccination conditions were analyzed by limma and edgeR, and a total of 613 DEGs were found, of which 304 DEGs were up-regulated and 309 DEGs were down-regulated in the vaccination group. These results are shown in heatmap which is a simple yet effective way to compare the content of multiple major gene lists (Fig. 3A-B).

**Fig. 3.**
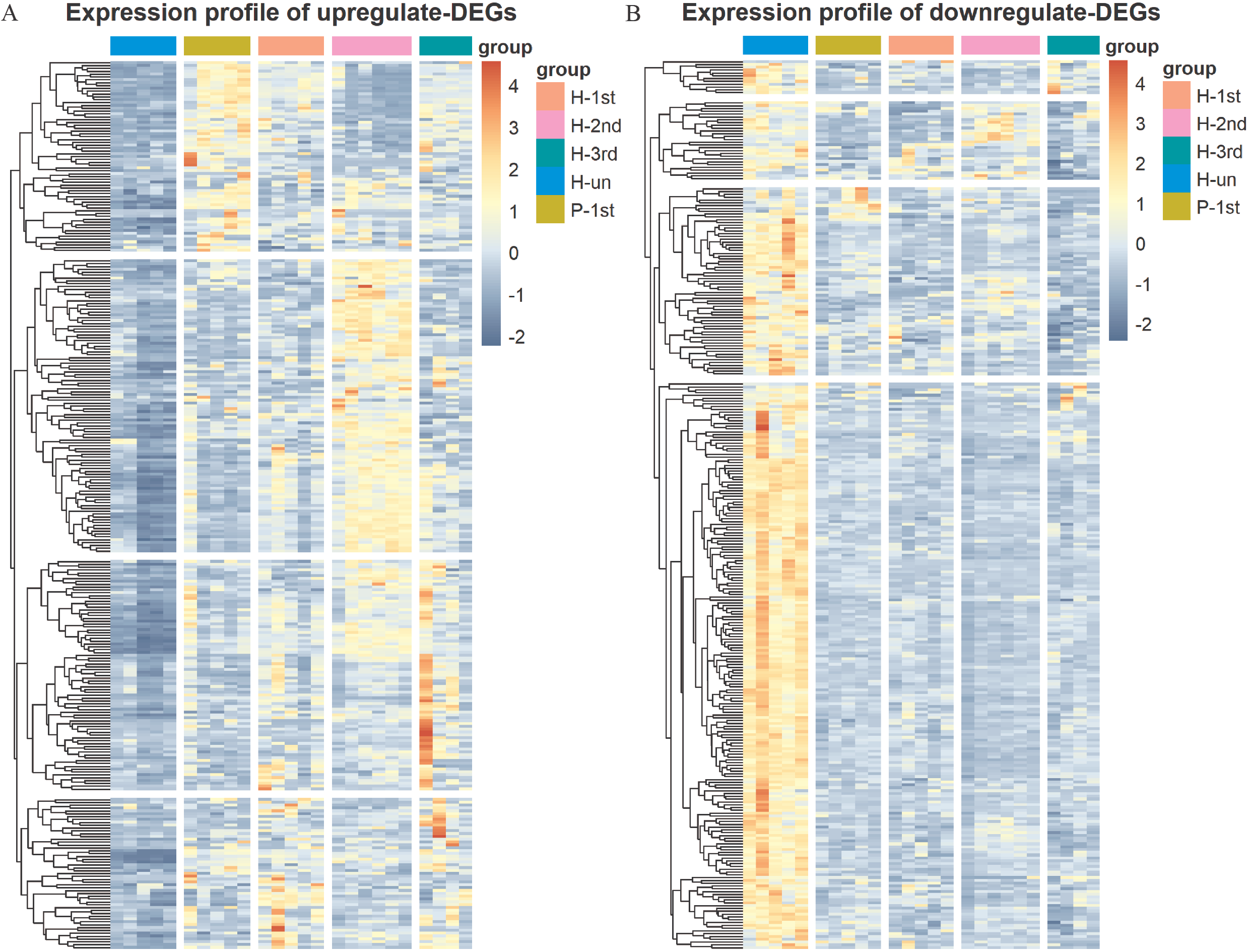
Hierarchical cluster analysis of DEGs in five groups. (A) The expression profile of 304 up-regulate DEGs. (B) The expression profile of 309 down-regulate DEGs. Each row represents mRNA and each column represents a sample. Orange indicates higher expression and blue indicates low expression in vaccination groups as compared with that in H-un group.

### Prioritization of DEGs by PPI network analysis

The PPI network of these up-regulated and down-regulated DEGs were established using STRING database, and the node and edge relationships were imported into Cytoscape software for visualization, and then the DC evaluation was completed. The degree results of 304 up-regulated DEGs and 309 down-regulated DEGs were demonstrated by the size of the point and the thickness of the edge (Fig. 4A and 5A). The first 61 up-regulated genes and 76 down-regulated genes were selected as characteristic genes (Fig. 4B and 5B) as a focus for subsequent examination. Using MCODE’s k-core decomposition function, a variety of sub-networks were obtained, and two new networks were created, in which 23 up-regulated and 35 down-regulated differentially expressed genes were retained, respectively (Fig. 4C and 5C). The results of DC and MCODE of up-regulated and down-regulated characteristic DEGs analysis are summarized in Tab. 1 and Tab. 2. The differential expression of these characteristic DEGs in the four vaccination groups is shown in the heatmap (Fig. 4D and 5D).

**Fig. 4.**
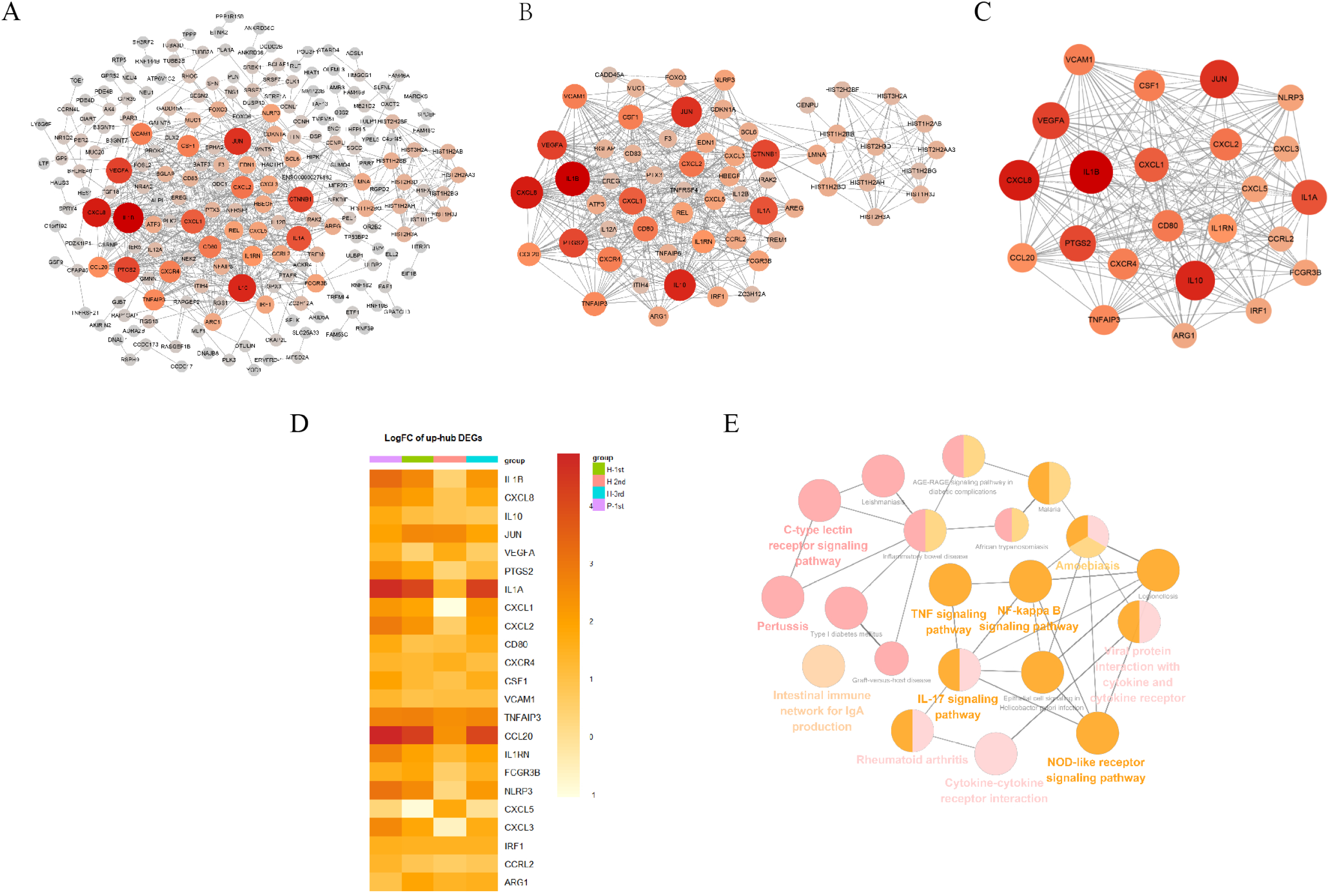
PPI networks of the up-regulated DEGs in vaccination groups. (A) The PPI network of all up-regulated DEGs, consisting of 304 nodes. (B) PPI network extracted from A by Degree Centrality, consisting of 61 nodes. (C) The PPI network extracted from B by MCODE, consisting of 32 nodes. PPI, protein-protein interaction; DEGs, differentially expressed genes. (D) The differential expression of 23 characteristic up-regulated DEGs in the four vaccination groups. Each row represents mRNA and each column represents a group. (E) A total of 35 KEGG pathways of the 23 characteristic up-regulated DEGs.

**Fig. 5.**
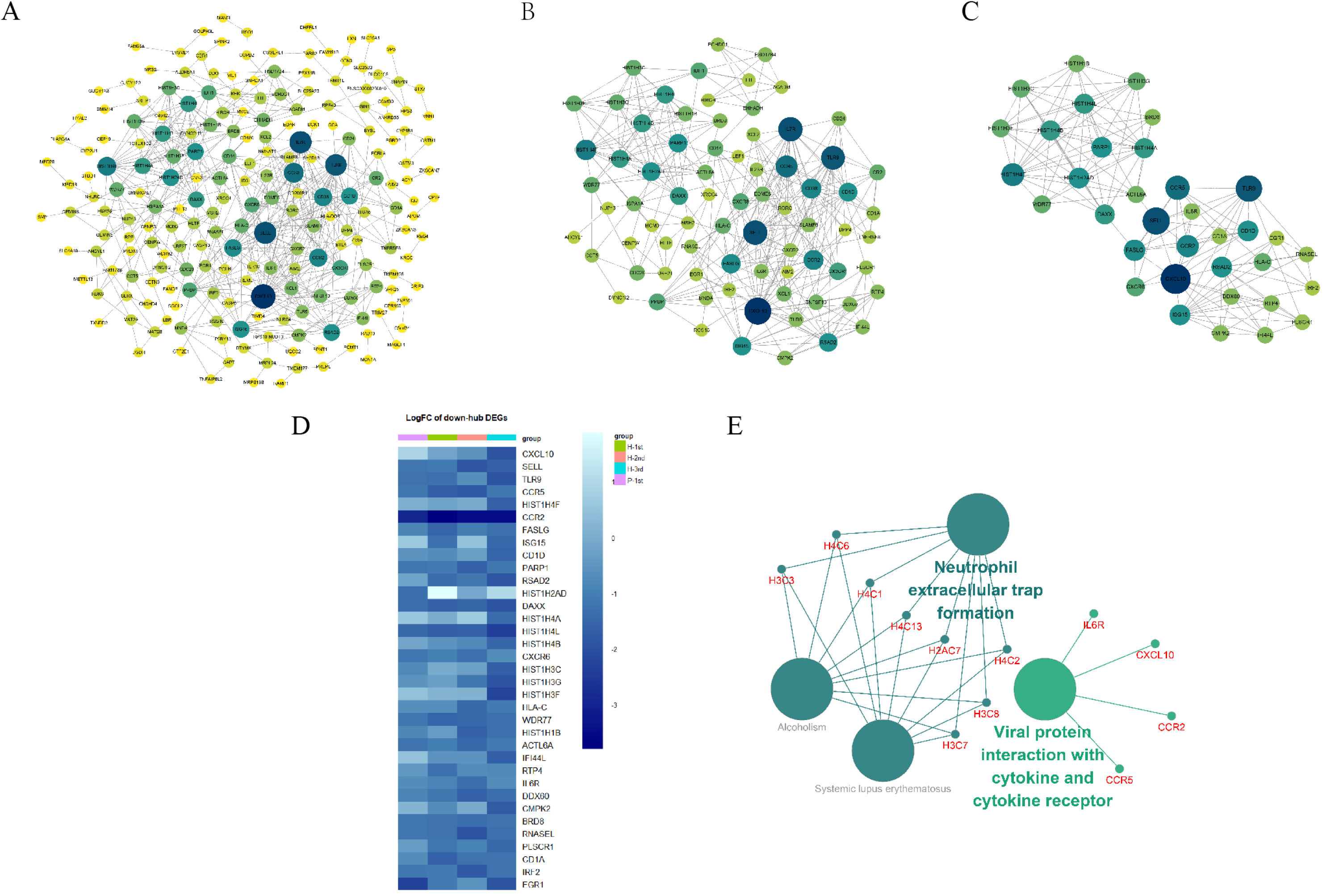
PPI networks of the down-regulated DEGs in vaccination groups. (A) The PPI network of all down-regulated DEGs, consisting of 309 nodes. (B) PPI network extracted from A by Degree Centrality, consisting of 76 nodes. (C) The PPI network extracted from B by MCODE, consisting of 35 nodes. PPI, protein-protein interaction; DEGs, differentially expressed genes. (D) The differential expression of 35 characteristic down-regulated DEGs in the four vaccination groups. Each row represents mRNA and each column represents a group. (E) A total of four KEGG pathways of the 35 characteristic down-regulated DEGs.

**Tab. 1.** Results of Degree Centrality and MCODE analysis of up-regulated characteristic genes.

| Name | Betweenness | Closeness | Degree | MCODE_Score | MCODE_Clusters |
| --- | --- | --- | --- | --- | --- |
| IL1B | 3002.72 | 2.71E-03 | 50 | 12.53 | Cluster 0 |
| CXCL8 | 1739.84 | 2.57E-03 | 45 | 12.53 | Cluster 0 |
| IL10 | 1238.76 | 2.50E-03 | 43 | 12.53 | Cluster 0 |
| JUN | 6024.51 | 2.78E-03 | 41 | 10.76 | Cluster 0 |
| VEGFA | 2147.79 | 2.53E-03 | 38 | 11.86 | Cluster 0 |
| PTGS2 | 1254.93 | 2.50E-03 | 37 | 12.53 | Cluster 0 |
| IL1A | 339.38 | 2.32E-03 | 35 | 12.53 | Cluster 0 |
| CXCL1 | 266.96 | 2.28E-03 | 33 | 12.53 | Cluster 0 |
| CXCL2 | 232.80 | 2.27E-03 | 29 | 12.53 | Cluster 0 |
| CD80 | 483.12 | 2.30E-03 | 28 | 12.88 | Cluster 0 |
| CXCR4 | 1655.69 | 2.38E-03 | 27 | 12.68 | Cluster 0 |
| CSF1 | 206.22 | 2.28E-03 | 27 | 12.53 | Cluster 0 |
| VCAM1 | 1170.73 | 2.34E-03 | 27 | 12.53 | Cluster 0 |
| TNFAIP3 | 1177.21 | 2.25E-03 | 25 | 10.40 | Cluster 0 |
| CCL20 | 123.02 | 2.12E-03 | 24 | 12.68 | Cluster 0 |
| IL1RN | 394.19 | 2.12E-03 | 23 | 13.00 | Cluster 0 |
| FCGR3B | 701.57 | 2.07E-03 | 18 | 11.00 | Cluster 0 |
| NLRP3 | 73.42 | 2.22E-03 | 18 | 11.30 | Cluster 0 |
| CXCL5 | 8.48 | 2.08E-03 | 18 | 12.43 | Cluster 0 |
| CXCL3 | 19.52 | 2.18E-03 | 17 | 12.88 | Cluster 0 |
| IRF1 | 28.39 | 2.15E-03 | 17 | 11.20 | Cluster 0 |
| CCRL2 | 48.71 | 2.02E-03 | 16 | 9.85 | Cluster 0 |
| ARG1 | 35.08 | 2.06E-03 | 16 | 13.00 | Cluster 0 |

**Tab. 2.** Results of Degree Centrality and MCODE analysis of down-regulated characteristic genes.

| Name | Betweenness | Closeness | Degree | MCODE_Score | MCODE_Clusters |
| --- | --- | --- | --- | --- | --- |
| CXCL10 | 4296.79 | 1.75E-03 | 27 | 6.61 | Cluster 0 |
| SELL | 3279.56 | 1.69E-03 | 23 | 6.61 | Cluster 0 |
| TLR9 | 2323.38 | 1.63E-03 | 22 | 5.79 | Cluster 0 |
| CCR5 | 2158.46 | 1.69E-03 | 19 | 6.61 | Cluster 0 |
| HIST1H4F | 2545.71 | 1.60E-03 | 17 | 8.00 | Cluster 0 |
| CCR2 | 1550.35 | 1.63E-03 | 16 | 6.61 | Cluster 0 |
| FASLG | 1833.84 | 1.62E-03 | 15 | 5.79 | Cluster 0 |
| ISG15 | 2379.69 | 1.53E-03 | 14 | 6.00 | Cluster 0 |
| CD1D | 212.26 | 1.56E-03 | 14 | 6.61 | Cluster 0 |
| PARP1 | 4184.37 | 1.64E-03 | 14 | 5.00 | Cluster 0 |
| RSAD2 | 591.91 | 1.50E-03 | 14 | 6.00 | Cluster 0 |
| HIST1H2AD | 1472.85 | 1.54E-03 | 14 | 8.00 | Cluster 0 |
| DAXX | 3097.22 | 1.64E-03 | 13 | 8.00 | Cluster 0 |
| HIST1H4A | 652.74 | 1.56E-03 | 13 | 8.00 | Cluster 0 |
| HIST1H4L | 652.74 | 1.56E-03 | 13 | 8.00 | Cluster 0 |
| HIST1H4B | 652.74 | 1.56E-03 | 13 | 8.00 | Cluster 0 |
| CXCR6 | 123.91 | 1.48E-03 | 11 | 5.00 | Cluster 0 |
| HIST1H3C | 396.62 | 1.43E-03 | 10 | 8.00 | Cluster 0 |
| HIST1H3G | 396.62 | 1.43E-03 | 10 | 8.00 | Cluster 0 |
| HIST1H3F | 396.62 | 1.43E-03 | 10 | 8.00 | Cluster 0 |
| HLA-C | 1079.68 | 1.48E-03 | 10 | 5.00 | Cluster 0 |
| WDR77 | 1201.87 | 1.47E-03 | 10 | 6.00 | Cluster 0 |
| HIST1H1B | 140.12 | 1.40E-03 | 9 | 7.00 | Cluster 0 |
| ACTL6A | 2228.69 | 1.58E-03 | 9 | 5.79 | Cluster 0 |
| IFI44L | 56.10 | 1.44E-03 | 8 | 6.00 | Cluster 0 |
| RTP4 | 56.10 | 1.44E-03 | 8 | 6.00 | Cluster 0 |
| IL6R | 68.42 | 1.49E-03 | 8 | 5.00 | Cluster 0 |
| DDX60 | 2385.37 | 1.48E-03 | 8 | 6.00 | Cluster 0 |
| CMPK2 | 1034.42 | 1.47E-03 | 8 | 6.00 | Cluster 0 |
| BRD8 | 3952.01 | 1.42E-03 | 8 | 6.00 | Cluster 0 |
| RNASEL | 653.95 | 1.45E-03 | 7 | 5.00 | Cluster 0 |
| PLSCR1 | 767.73 | 1.38E-03 | 7 | 5.00 | Cluster 0 |
| CD1A | 38.07 | 1.39E-03 | 7 | 5.00 | Cluster 0 |
| IRF2 | 370.00 | 1.32E-03 | 6 | 5.00 | Cluster 0 |
| EGR1 | 258.74 | 1.38E-03 | 6 | 5.00 | Cluster 0 |

### KEGG pathway enrichment analyses of characteristic genes

The KEGG term enrichment analyses were performed using CluGO. There were 19 KEGG pathways with P<0.05 (including Bonferroni adjustment) identified within the analysis of the 23 characteristic up-regulated genes, including TNF signaling pathway, IL-17 signaling pathway, Viral protein interaction with cytokine and cytokine receptor, Cytokine-cytokine receptor interaction, Pertussis, NF-kappa B signaling pathway and other pathways (Fig. 4E). In these pathways, genes that play important roles include chemokine, interleukin, macrophage colony-stimulating factor, interferon regulatory factor 1 and tumor necrosis factor alpha-induced protein 3. There were four KEGG pathways with P<0.05 (including Bonferroni adjustment) identified within the analysis of the 35 characteristic down-regulated genes, including Systemic lupus erythematosus, Alcoholism, Neutrophil extracellular trap formation, Viral protein interaction with cytokine and cytokine receptor (Fig. 5E). Among these pathways, it was hypothesized that genes which serve a significant role include histone cluster, chemokine and interleukin-6 receptor subunit alpha (IL6R). Detailed enrichment information for the KEGG pathways is summarized in Tab. 3-4.

**Tab. 3.** KEGG pathway of the 23 characteristic up-regulated genes.

| ID | Term | P-value | Associated Genes Found |
| --- | --- | --- | --- |
| KEGG:04668 | TNF signaling pathway | 1.56E-17 | [CCL20, CSF1, CXCL1, CXCL2, CXCL3, CXCL5, IL1B, IRF1, JUN, PTGS2, TNFAIP3, VCAM1] |
| KEGG:04657 | IL-17 signaling pathway | 1.58E-14 | [CCL20, CXCL1, CXCL2, CXCL3, CXCL5, CXCL8, IL1B, JUN, PTGS2, TNFAIP3] |
| KEGG:04061 | Viral protein interaction with cytokine and cytokine receptor | 2.03E-12 | [CCL20, CSF1, CXCL1, CXCL2, CXCL3, CXCL5, CXCL8, CXCR4, IL10] |
| KEGG:04060 | Cytokine-cytokine receptor interaction | 2.06E-12 | [CCL20, CSF1, CXCL1, CXCL2, CXCL3, CXCL5, CXCL8, CXCR4, IL10, IL1A, IL1B, IL1RN] |
| KEGG:05133 | Pertussis | 1.19E-11 | [CXCL5, CXCL8, IL10, IL1A, IL1B, IRF1, JUN, NLRP3] |
| KEGG:04064 | NF-kappa B signaling pathway | 1.56E-10 | [CXCL1, CXCL2, CXCL3, CXCL8, IL1B, PTGS2, TNFAIP3, VCAM1] |
| KEGG:05146 | Amoebiasis | 5.99E-09 | [ARG1, CXCL1, CXCL2, CXCL3, CXCL8, IL10, IL1B] |
| KEGG:04621 | NOD-like receptor signaling pathway | 1.31E-08 | [CXCL1, CXCL2, CXCL3, CXCL8, IL1B, JUN, NLRP3, TNFAIP3] |
| KEGG:05140 | Leishmaniasis | 4.02E-08 | [FCGR3B, IL10, IL1A, IL1B, JUN, PTGS2] |
| KEGG:04933 | AGE-RAGE signaling pathway in diabetic complications | 1.95E-07 | [CXCL8, IL1A, IL1B, JUN, VCAM1, VEGFA] |
| KEGG:04625 | C-type lectin receptor signaling pathway | 2.46E-07 | [IL10, IL1B, IRF1, JUN, NLRP3, PTGS2] |
| KEGG:05134 | Legionellosis | 3.48E-07 | [CXCL1, CXCL2, CXCL3, CXCL8, IL1B] |
| KEGG:05120 | Epithelial cell signaling in Helicobacter pylori infection | 9.84E-07 | [CXCL1, CXCL2, CXCL3, CXCL8, JUN] |
| KEGG:05144 | Malaria | 8.69E-06 | [CXCL8, IL10, IL1B, VCAM1] |
| KEGG:05321 | Inflammatory bowel disease | 2.49E-05 | [IL10, IL1A, IL1B, JUN] |
| KEGG:05143 | African trypanosomiasis | 1.28E-04 | [IL10, IL1B, VCAM1] |
| KEGG:05332 | Graft-versus-host disease | 1.87E-04 | [CD80, IL1A, IL1B] |
| KEGG:04940 | Type I diabetes mellitus | 2.01E-04 | [CD80, IL1A, IL1B] |
| KEGG:04672 | Intestinal immune<br>network for IgA<br>production | 2.96E-04 | [CD80, CXCR4, IL10] |

**Tab. 4.**
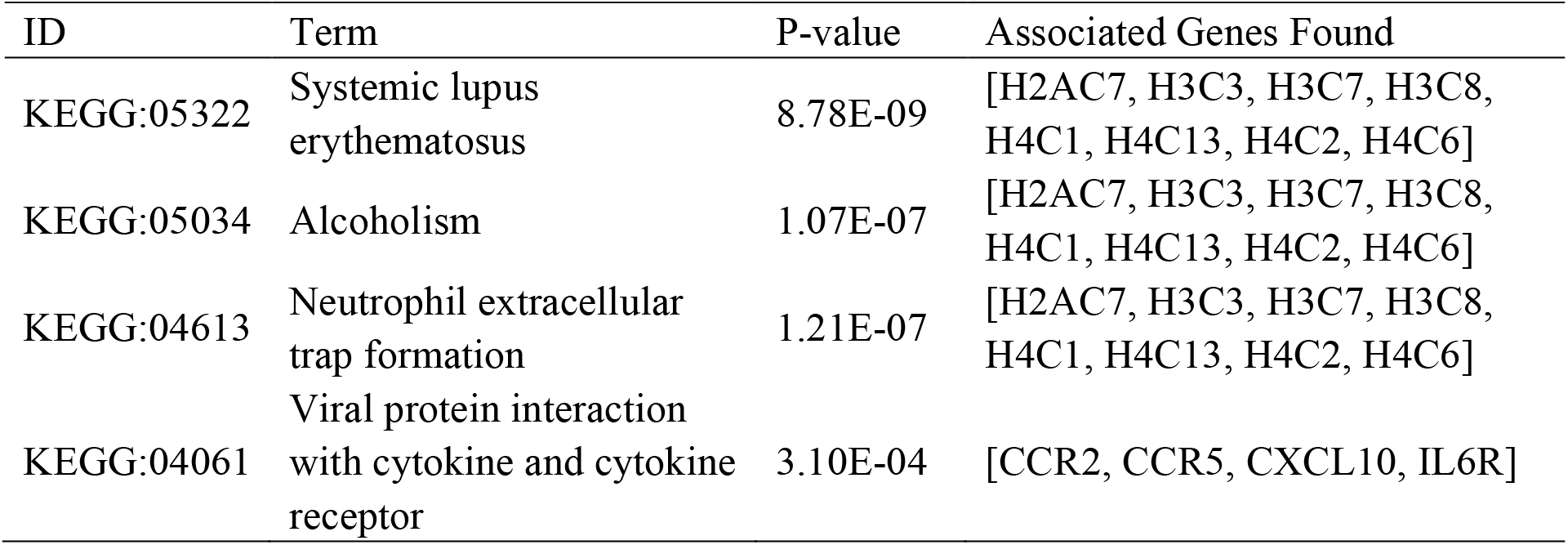
KEGG pathways of the 35 characteristic down-regulated genes.

### WGCNA: Identify important feature modules related to sample features

To explore the impact of different vaccine doses and SARS-CoV-2 infection status on the transcriptome, we made an analysis of the transcriptome using WGCNA. In this study, the power of β = 5(scale-free R 2 = 0.85) was selected as the soft-thresholding parameter to ensure a scale-free network (Fig. 6A-B). A total of 37 modules were identified through hierarchical clustering, and a dendrogram of all DEGs was clustered based on a dissimilarity measure (1-TOM) (Fig. 6C-D). The correlation between the sample characteristics and the co-expression module is shown in Fig. 6E, where the Brown (eigengene value = 0.69) module is significantly positively correlated with the vaccine dose. The darkorange (eigengene value = 0.91), yellow (eigengene value = 0.81) and steelblue (eigengene value = 0.71) modules are significantly correlated with the previous SARS-CoV-2 infection status (Fig. 6F-I).

**Fig. 6.**
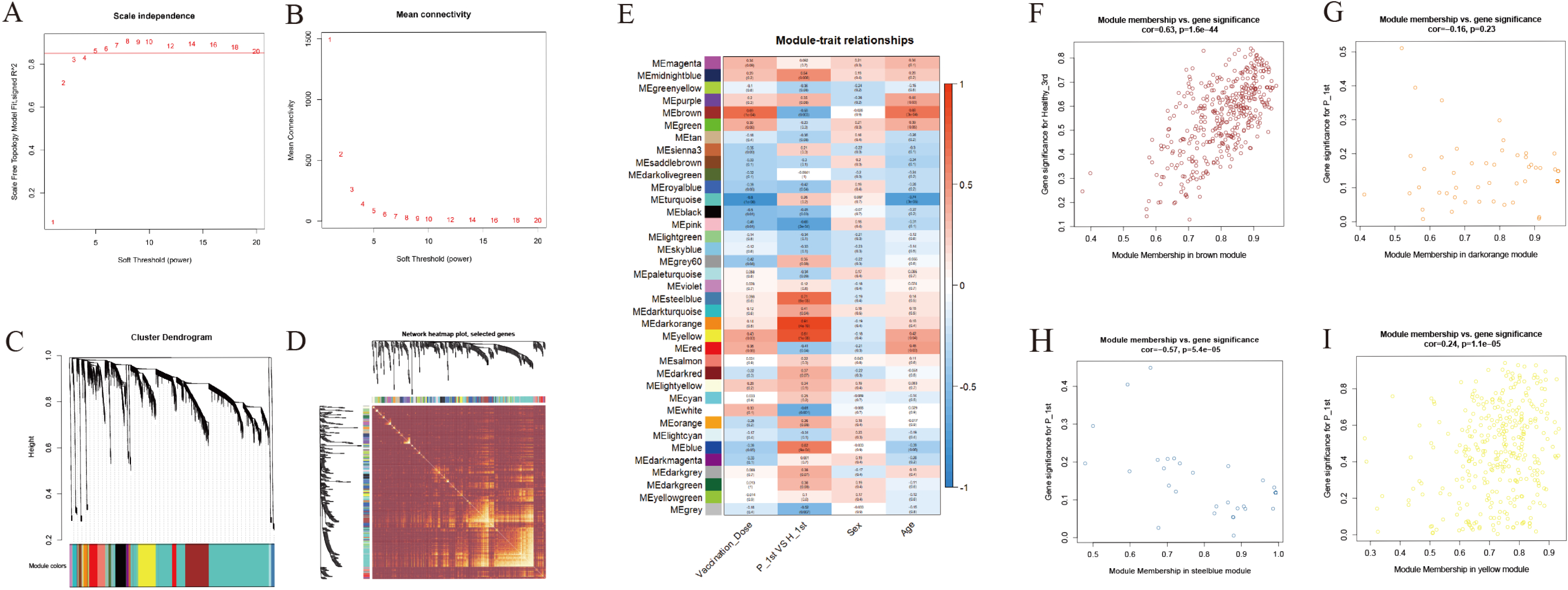
WGCNA of the PBMCs transcriptome. (A-B) Analysis of network topology for various soft-thresholding powers. The left panel shows the scale-free fit index (y-axis) as a function of the soft-thresholding power (x-axis). The right panel displays the mean connectivity (degree, y-axis) as a function of the soft-thresholding power (x-axis). (C) Clustering dendrogram of genes, with dissimilarity based on topological overlap, together with assigned module colors. (D) Heatmap depicts the Topological Overlap Matrix (TOM) of genes selected for weighted co-expression network analysis. Light color represents lower overlap and red represents higher overlap. (E) Module-trait associations: Each row corresponds to a module eigengene and each column to a trait. Each cell contains the corresponding correlation and p-value. (F-I) Scatter diagram for MM vs GS in the brown, darkorange, yellow and steelblue module.

### Analysis of important modules and identification of genes

In order to analyze the network in the corresponding modules and determine the importance and function of genes, we combined CytoHubba and GeneMANIA in Cytoscape to analyze and identify the hub genes of the above four modules. The functions of the hub gene of the brown module mainly focus on activation of innate immune response, response to interleukin-1, and cellular response to tumor necrosis factor, etc. The hub genes of darkorange module act on regulation of tyrosine phosphorylation of STAT protein, tyrosine phosphorylation of STAT protein, pattern specification process, postsynaptic neurotransmitter receptor activity, neurotransmitter receptor activity pathway. The hub gene of the yellow module performs adaptive immune response, regulation of T cell activation, and B cell mediated immunity, etc. The hub gene of steelblue module can participate in humoral immune response, detection of biotic stimulus, antimicrobial humoral response, glycosaminoglycan binding, peptidyl-arginine modification and other pathways (Fig. 7A-D).

**Fig. 7.**
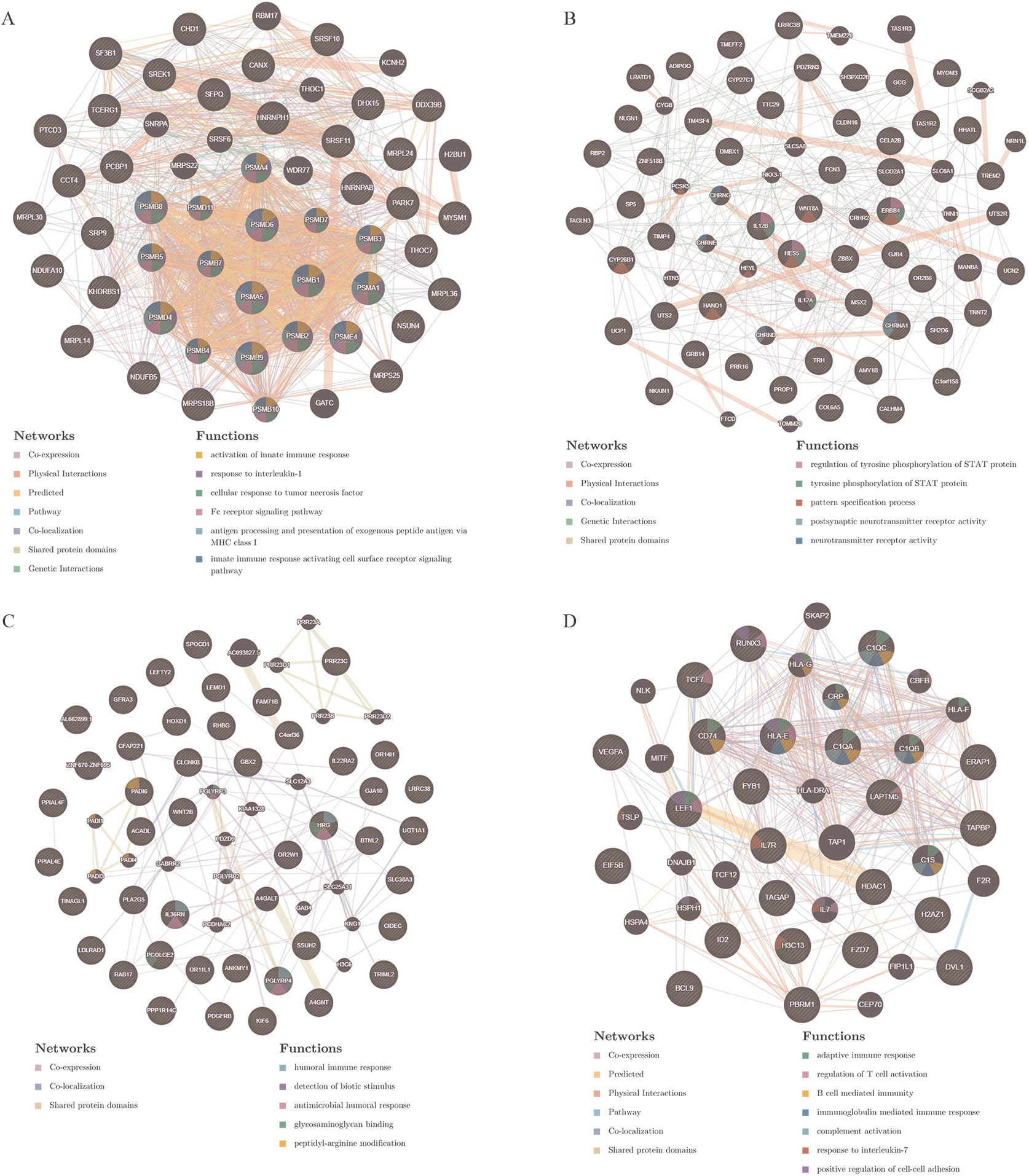
PPI network construction. The PPI network of brown (A), darkorange (B), yellow (C) and steelblue (D) module. Different colors in each node represent different functions as indicated. Function of genes in dark node was not revealed through the analysis. Different colors in each edge represent different connection.

## Discussion

The SARS-CoV-2 pandemic poses an imminent threat to humans, and the development and application of vaccines have brought hope to the effective prevention and control of the epidemic. In the field of rapid development and promotion of vaccines, the real data of vaccinated persons is crucial to the management and control of COVID-19 by government agencies and public health departments. At present, there is no research on the transcriptome response of COVID-19 recovered individuals and SARS-CoV-2-naïve individuals after vaccination with COVID-19 inactivated vaccine. In this study, we systematically revealed the molecular immune response and response mechanism of the vaccinated individuals to the inactivated vaccine.

Compared with the unvaccinated population, the up-regulated characteristic genes with substantial transcriptional changes in the vaccinated groups included IL1B, CXCL8, IL10, JUN, VEGFA, PTGS2, IL1A, CXCL1, CXCL2, CD80, etc. They participate in the activation of many innate immune pathways, including TNF signaling pathway, IL-17 signaling pathway, Viral protein interaction with cytokine and cytokine receptor, Cytokine-cytokine receptor interaction, Pertussis, NF-kappa B signaling pathway, etc. These transcriptional responses are highly consistent with the transcriptional regulation induced by inactivated vaccines of other pathogens. In research on inactivated influenza vaccines, the expression of IL-17 and NF-κB pathway genes and related innate immune activation have been observed(12); Rubella vaccine significantly affects the host’s antigen presentation and innate/inflammatory genome(22); Hantavax vaccine can mediate Th17 cell differentiation, antigen processing and presentation, NF-kappa B signalling pathway, phenylalanine metabolism, phagosome, Fc gamma R-mediated phagocytosis(11). In the study of BNT162b2 mRNA vaccine, it was found that vaccination not only stimulated antiviral and interferon response modules, but also induced a broader innate immune response such as Toll-like receptor signalling, monocyte and neutrophil modules(23). The above research results emphasize the key role of innate immune response in vaccination.

The characteristic genes that were significantly down-regulated in the vaccinated group included Histone cluster, chemokine and IL6R that participated in Systemic lupus erythematosus, Alcoholism, Neutrophil extracellular trap formation, Viral protein interaction with cytokine and cytokine receptor pathways. In a systems biology analysis of PBMC transcriptome data from COVID-19 patients, it was also found that key gene modules are not only involved in infection-related pathways, but also significantly enriched in Systemic lupus erythematosus and Alcoholism(24). This coincidental consistency suggests that the role of these two pathways in COVID-19 needs to be further explored. In a study on Live-Attenuated *Francisella tularensis* vaccine, it was found that while a variety of immune pathways were activated early, some innate immune signaling pathways were inhibited, especially cytokine-cytokine receptor interactions.

This result is related to the host escape mechanism of *Francisella tularensis*(25). Recent investigations suggested that in addition to the virus itself, dysregulated host immune response to SARS-CoV-2 may also contribute to the pathogenesis of COVID-19(26–28). The inhibition of Neutrophil extracellular trap formation and Viral protein interaction with cytokine and cytokine receptor pathway caused by COVID-19 inactivated vaccine may also be related to the immune escape of SARS-CoV-2.

For the SARS-CoV-2 naïve individuals, the determination of the number of doses is critical to the large-scale vaccination and production requirements of the vaccine. In the early days, a two-dose vaccination program was implemented in the SARS-CoV-2 naïve individuals, with an interval of 14 days between the two doses. Now we are trying to carry out a third booster vaccine half a year after the second vaccine. Our research data found that although the first dose of vaccine played a partial regulatory role at the transcriptome level, the positive rate of IgG antibodies in serum samples was very low. After the second and third doses of vaccine, the IgG antibody positive rate reached 100%. WGCNA analysis shows that with the increase of vaccine doses, it is significantly related to the brown modules involved in activation of innate immune response, response to interleukin-1, cellular response to tumor necrosis factor, Fc receptor signaling pathway, antigen processing and presentation of exogenous peptide antigen via MHC class I and innate immune response activating cell surface receptor signaling pathway. These functions are mainly performed by hub genes related to the 20S proteasome subunit, including PSMA1, PSMA4, PSMA5, PSMB1, PSMB10, PSMB2, PSMB3, PSMB4, PSMB5, PSMB7, PSMB8, PSMB9, PSMD11, PSMD4, PSMD6, PSMD7, PSME4. The proteasome is a multicatalytic proteinase complex with a highly ordered ring-shaped 20S core structure. The 20S proteasome is formed by the 14 α subunits and the 14 β subunits, in which β1 (coded by the PSMB6 gene), β2 (coded by PSMB7 gene), and β5 (coded by PSMB5 gene) subunits possess protease activities(29). It is the catalytic core of the 26S proteasome and is distributed throughout eukaryotic cells at a high concentration(30). An essential function of a modified proteasome is the processing of class I MHC peptides. It can continue to decompose intracellular proteins that may appear in the process of viral infection, and produce antigen peptides, which can be combined with MHC1 molecules in the endoplasmic reticulum and transport the complex to the cell surface to play a role in antigen presentation(31). Obviously, the function of the proteasome in the immune system is significant, and the third booster vaccine is beneficial and harmless. A pivotal question in the process of vaccination promotion is whether individuals who have previously been infected with SARS-CoV-2 need to be vaccinated with multiple doses, because these people have developed a primary immune response to the virus during natural infection(32–34), so a single dose of vaccine may be sufficient enhance immune response. This issue is particularly important in environments where vaccine supply is limited and vaccine deployment is challenging. Judging from the results of the IgG antibody response of the serum samples, the recovered patients have already had a similar effect to the two or three doses of vaccines given to uninfected individuals after receiving one dose of vaccine. Co-expression analysis of transcriptome data showed that darkorange, yellow, and steelblue modules were significantly related to SARS-CoV-2 infection status in people who received one dose of vaccine. The hub genes of these modules perform many adaptive immune regulation functions, including cellular immune responses such as the activation and regulation of T cells, and humoral immune responses mediated by B cells and antibodies. It suggests that a single dose of the inactivated vaccine elicits a strong immune response in population vaccinated 18 months after recovering from COVID-19. In some study of mRNA vaccines on SARS-CoV-2 naïve and recovered individuals, it was found that after a single dose of vaccine, recovered individuals can produce a strong immune response, including serum antibody response, B cell response, and transcriptome changes(14, 35, 36). The current results show that in the case of limited vaccine supply, we can consider reducing the number of vaccines for SARS-CoV-2 recovered individuals. However, in view of the relatively short duration of this study, future studies will be necessary to evaluate the impact of multiple doses of vaccine on the long-term immune response of recovered individuals. Our data provides quantitative and global insights into the dynamic changes of immune-related genes, reveals the differential expression of cytokines and inflammatory factors, as well as the regulation of antigen presentation and innate immune pathways, and helps us explain the effects of molecular immune mechanism induced by inactivated vaccines. The transcriptome characteristics of individuals with different SARS-CoV-2 infection and different doses may provide a new reference for the determination of vaccination strategies in the future. However, the results of this study rely on bioinformatics analysis, the sample size is small, and the long-term evaluation of the inoculation effect is lacking. We intend to further expand the sample size and conduct long-term follow-up to explore comprehensive insights into the inactivated COVID-19 vaccine.

## Contributors

X.L.J., and Y.W.Z. conceived the study. Y.W.Z. devised the statistical methodology; C.B.L. and L.F.L. recruited participants and collected samples; Y.W.Z., X.Y.G., L.F.L., M.X.Y, and B. P. did the formal analysis; X.L.J., Y.W.Z., and Z.Q.K. were project administrators; X.M.Z., and Q.D. curated and validated the data; X.Y.G., X.Y.T., and Y.F.X. did the literature review. X.L.J. acquired the funding; Y.W.Z., and X.Y.G. wrote the original draft of the manuscript; X.L.J. reviewed and edited the manuscript; X.L.J. and Y.W.Z. have shared responsibility for the decision to submit for publication. All authors reviewed and approved the final manuscript.

## Declaration of interests

The authors declare that they have no competing interests.

## Role of the funding source

This work was supported by the Major Scientific and Technological Innovation Project in Shandong Province(grant numbers: 2020SFXGFY02-1).

## Data sharing

The RNA-seq data from this study will be uploaded in GEO before publishing the manuscript, and be made available to others with publication.

